# DeepLC can predict retention times for peptides that carry as-yet unseen modifications

**DOI:** 10.1101/2020.03.28.013003

**Authors:** Robbin Bouwmeester, Ralf Gabriels, Niels Hulstaert, Lennart Martens, Sven Degroeve

## Abstract

The inclusion of peptide retention time prediction promises to remove peptide identification ambiguity in complex LC-MS identification workflows. However, due to the way peptides are encoded in current prediction models, accurate retention times cannot be predicted for modified peptides. This is especially problematic for fledgling open modification searches, which will benefit from accurate retention time prediction for modified peptides to reduce identification ambiguity. We here therefore present DeepLC, a novel deep learning peptide retention time predictor utilizing a new peptide encoding based on atomic composition that allows the retention time of (previously unseen) modified peptides to be predicted accurately. We show that DeepLC performs similarly to current state-of-the-art approaches for unmodified peptides, and, more importantly, accurately predicts retention times for modifications not seen during training. Moreover, we show that DeepLC’s ability to predict retention times for any modification enables potentially incorrect identifications to be flagged in an open modification search of CD8-positive T-cell proteome data. DeepLC is available under the permissive Apache 2.0 open source license and comes with a user-friendly graphical user interface, as well as a Python package on PyPI, Bioconda, and BioContainers for effortless workflow integration.

## Introduction

Liquid Chromatography (LC) plays a critical role in Mass Spectrometry (MS) analysis of bottom-up proteomics^1^. By separating peptides based on their physicochemical properties in the LC step, the complexity of the sample presented to the MS instrument is greatly reduced. This reduction means that there is less ionization competition, improved sensitivity for data dependent/independent analysis, and reduced chimericity in fragmentation spectra (MS^2^) ^2,3^. In addition to these benefits, the retention time measurement itself provides an additional dimension of information to interpret the signals generated by a peptide^4^. In order to interpret these acquired signals, they need to be matched with earlier observations of the same peptides or with a prediction of the signal.

However, the process by which a peptide is retained or eluted is not fully understood yet^5^, which means that libraries with previously observed retention times are often used to match to newly acquired signals^6^. However, these libraries are often incomplete and moreover are non-transferable between experimental setups without calibration. In order to fill this knowledge gap, researchers have therefore used models to predict retention times for previously unobserved peptides^4^.

Many of the first methods for peptide retention time prediction relied on simulation models based on physicochemical knowledge^7^. However, most modern approaches use data-driven methods such as machine learning (ML) or deep learning (DL) algorithms to train a predictive model^8–12^. In such models, the mapping between the peptide sequence (or features derived from this sequence) and the LC retention time apex is learned from empirical examples. After training, these models can be used to generate predictions for unobserved peptides.

Such retention time prediction models have already been successfully applied for various tasks in proteomics analysis workflows, e.g. to improve identification confidence^13,14^, to design more efficient experiments^15^, and to identify chimeric fragmentation spectra^16^. Most recently, these retention time prediction models have been used in combination with fragment peak intensity prediction models to generate comprehensive, *in silico* chromatogram libraries for Data Independent Acquisition (DIA) identification, effectively replacing and surpassing more limited, empirically derived Data Dependent Acquisition (DDA) spectral libraries^17–19^.

In keeping with the general trend in ML, there has been a switch from classical ML to DL in newly developed retention time predictors. This switch was mainly driven by recent innovations in the field of DL and the large amount of peptide retention time data that has become available. The types of architectures proposed by state-of-the-art DL retention time models include capsule convolutional neural networks (CNNs) in DeepRT(+)^11^, NN with long short-term memory (LSTM) layers as used by Guan et al.^9^, and an encoder-decoder principle with gated recurrent units (GRU) in Prosit^10^. The architectures of these models either work with a CNN or recurrent architecture (e.g. LSTM or GRU units). CNN architectures slide a filter with a specified kernel size over the encoded peptide. In contrast, recurrent neural networks thread the sequence through a sequence of units.

However, all these models share the same peptide encoding method, in which amino acids and their corresponding positions are transformed into a one-hot amino acid encoding. This encoding takes the form of a matrix in which the presence or absence of each amino acid for each position in the peptide is represented by a one or a zero, respectively.

Unfortunately, this use of one-hot encoding of amino acids restricts the models’ applicability in some of the most interesting data analysis workflows, most notably in open modification searches (OMS), where the goal is to elucidate the modification landscape of the proteome^20–23^. These OMS workflows are gaining popularity in the field of proteomics as they make it possible to search for a large variety of peptide modifications simultaneously.

Unfortunately, current retention time prediction methods cannot be directly applied in OMS because of the vast amount of potential modifications^24^. With one-hot amino acid encoding, each potential modification needs to be represented by a binary feature indicating the presence of this modification. Additionally, sufficient training examples are required for each modification for the ML algorithm to learn the hidden impact of every one of these modifications on the peptide retention time.

We here solve this fundamental issue with DeepLC, our novel retention time predictor that is able to accurately predict the retention time for all peptides and their modifications, even when these modifications have not been seen during training. DeepLC achieves this by encoding peptides and modifications at the atomic composition level, allowing generalization of the patterns learned from the modifications seen during training.

## Results

The results section is split in two main parts. We first evaluate the performance of DeepLC on retention time prediction for unmodified peptides, in comparison with state-of-the-art tools. We then proceed to evaluate DeepLC on retention time prediction for modified peptides, which is an ability that is unique to DeepLC. We therefore rely on two distinct ways of evaluating DeepLC’s performance on these modified peptides: (i) evaluate DeepLC performance on unseen modifications, and (ii) a novel type of evaluation which leaves out unmodified amino acids, and then has DeepLC treat these as modified glycines. These evaluations show that DeepLC is not only competitive with state-of-the-art retention time prediction algorithms for unmodified peptides, but can also achieve similar performance for unseen modified peptides. Finally, we illustrate a practical use of this unique capability by showing DeepLC’s ability to flag false positive identifications in the results of an open modification search of a CD8-positive T-cell data set.

### Evaluation of DeepLC against the state-of-the-art on unmodified peptides

Our approach to model amino acids by their atomic composition provides accurate predictions of LC retention times for unmodified peptides, with similar performance to current state-of-the-art retention time prediction models DeepRT^11^, Prosit^10^, and Guan et al.^9^ that model amino acids directly. Note that the definition used here for “unmodified” can include the two very common artefactual modifications of carbamidomethylation of cysteine and oxidation of methionine.

DeepLC test set predictions for the three selected data sets are plotted in Figure 1. We observe very high prediction accuracy, with Pearson correlations larger than 0.99 for two of the data sets, and with the *HeLa HF* data set showing slightly worse performance (Pearson=0.984). For the latter, the LC gradient is significantly shorter than for the other two data sets, implying that retention times become less predictable, thus resulting in a slightly less performant model. Figure 1 also reveals a small but significant number of peptides with high prediction errors that do not follow the main trend. These are potentially wrong identifications or wrongly determined elution apexes. The same plots for the other seventeen data sets can be found in Supplementary Figures 1 and 2 where we make very similar observations to those in Figure 1.

**Figure 1:**
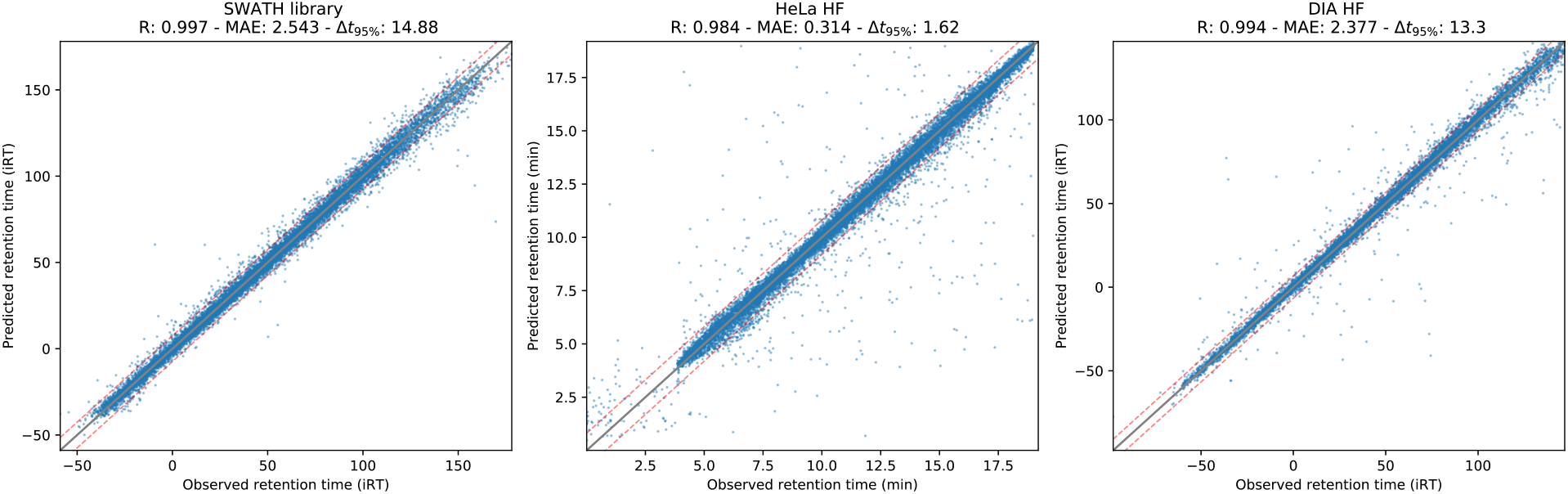
Scatter plot of predicted against observed on three of the largest data sets; *SWATH Library, HeLa HF*, and *DIA HF*.

Supplementary Table 1 summarizes the test set performance for all twenty data sets described in the data sets & evaluation section of the methods. The atomic composition encoding approach of DeepLC is able to learn accurate prediction models with high R values for all data sets. For most data sets DeepLC achieves an R above 0.98, with four data sets even achieving correlations above 0.995. This R value is highly comparable to the other models. Nearly all data sets were obtained with reverse phase columns, yet even though there are fewer data sets with HILIC and SCX, DeepLC also performs very well on these data sets with relative MAE errors below 1.5 %.

For the ∆*t*_95%_ metric the differences are somewhat more pronounced. Here we observe that DeepLC performs consistently worse than the other models. It is however unclear whether these differences should be accounted to the atomic composition encoding, a different DL architecture, a difference in train-validation-test split (note that for the other prediction models the manuscript did not mention the use of a validation set), or a combination of these. As we want to focus on the capability of DeepLC to predict retention times for modified peptides we leave this question open for further research.

The trained models are also highly transferrable between different data sets. This transferability is especially useful when applying models trained on larger data sets to smaller ones and application of the models without retraining. Only a simple calibration is required to transfer the predictions between LC setups. Supplementary Figure 3 shows that models that achieve high performance on a given data set also show high Spearman correlation when applied to different data sets.

DeepLC builds on a DL approach that greatly benefits from a large number of training peptides, and we can show that large data sets indeed do have a positive influence on the performance of DeepLC. The performance on each individual data set in relation to the number of training peptides is shown in Figure 2 and Supplementary Table 1. Data sets with a very small number of training peptides (< 10 000) tend to have a performance between 2 % and 4.5 % relative MAE. For medium sized data sets (> 10 000 and < 75 000 peptides), the performance can vary widely, with relative MAE’s ranging from 0.9 % to 4.5 %. For larger data sets (> 75 000 peptides) the performance tends to be below 2 % relative MAE.

**Figure 2:**
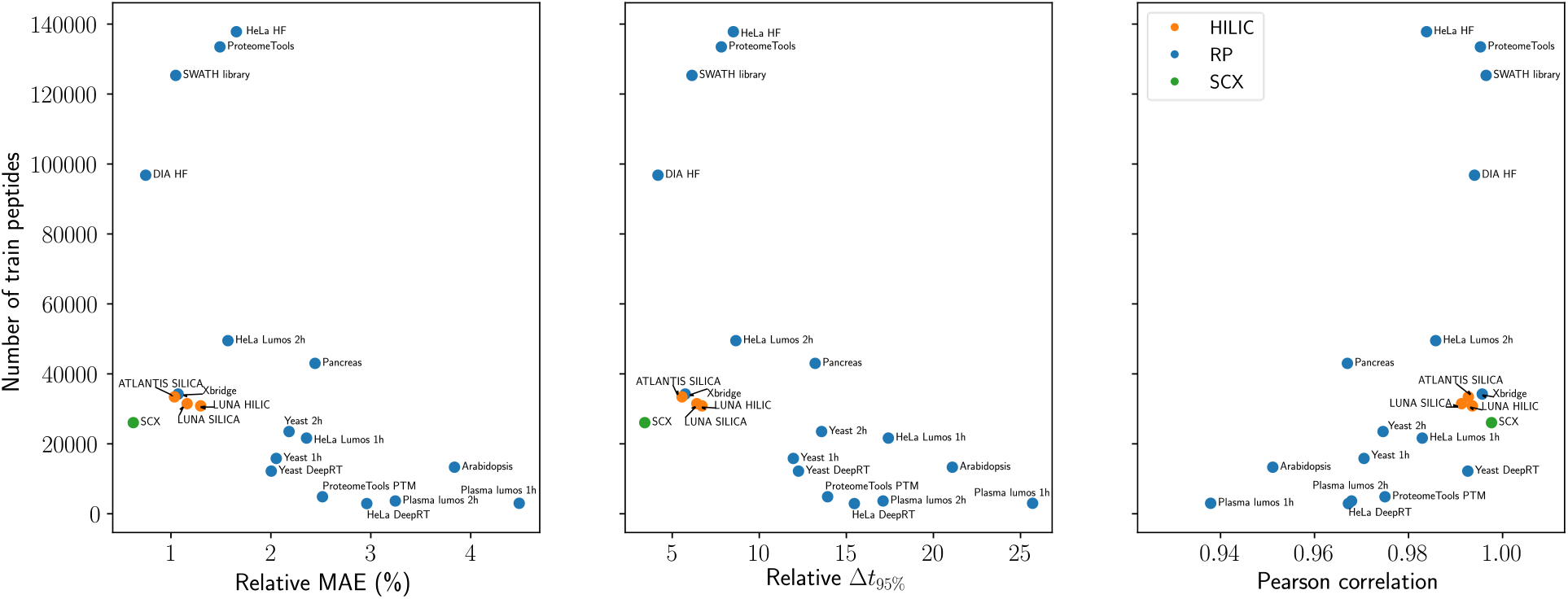
For the twenty data sets the number of training peptides (y-axis) is plotted against the relative MAE (left), relative Δ*t*_95%_ (middle) and Pearson correlation (right).

To further evaluate the relationship between the number of training peptides and prediction performance we computed learning curves for the three selected data sets (Figure 3). For each data set, learning curve evaluation was performed five times with different test set subsamples. These curves show a sharp improvement for the first four to five steps (comprising up to 50 percent of the total number of training peptides). Beyond these steps, prediction performance improves only linearly for the *SWATH library* and *HeLa HF*, while showing smaller improvements in the last step for *DIA HF*. Importantly, for two of these data sets the performance continues to improve right to the last step of the learning curve. This ability to continuously improve performance suggests that DeepLC, like most other DL approaches, is capable of fitting even more complex relations than classical ML when provided with sufficient data. The same observation of increasing performance for larger training sets can be made for the remaining seventeen data sets (Supplementary Figures 4 and 5), with fifteen data sets showing clear improvement for the last two steps of the learning curve.

**Figure 3:**
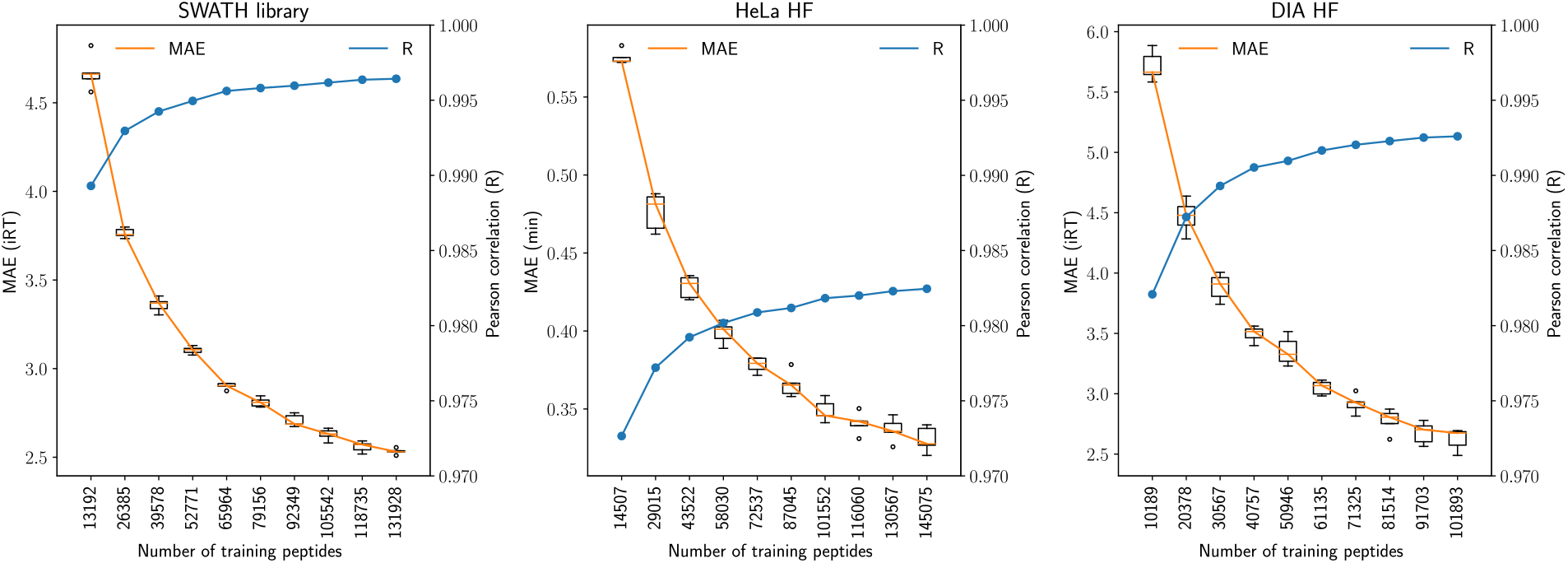
Learning curves for each of the three selected data sets. Prediction performances (R and MAE) for models trained on different training set sizes (x-axis) are computed for a fixed test set.

### Performance evaluation for modified peptides

DeepLC is able to generalize effectively for unmodified peptides as well as extending its accurate retention time predictions to modifications that were not included in the training set. We can thus show that the DeepLC models have not just learned the general shift in retention time caused by modifications, but also how this shift depends on the context of the modification in the peptide. These claims are supported by two separate evaluations in this section.

Prediction performance for modified peptides would ideally be evaluated on a large data set with a variety of modifications. Indeed, as shown in Figure 3, the full performance potential of DeepLC is achieved by the largest possible data set size. However, such large data sets with many modifications are currently not available in the public domain.

Instead, we here show DeepLC’s prediction performance for modified peptides on a recently published smaller data set (*ProteomeTools PTM*^25^), after the custom preprocessing workflow described in the methods section, comprising fourteen different modifications with known location in 4099 synthetic peptides. Furthermore, we introduce an evaluation procedure that allows the use of larger data sets based on the fact that any amino acid (apart from glycine) can be considered as a modified glycine.

We first evaluate DeepLC on all fourteen modifications in the *ProteomeTools PTM* data set. We trained and optimized fourteen DeepLC models, where each model only sees peptides that don’t contain a specific modification *M*. Each model is named after the modification it was not provided with, so model *M* is trained and optimized only on those peptides that do not contain modification *M*. Each model is then evaluated on the remaining peptides, which all do contain the modification *M* that was excluded during training. We created two test sets from these remaining peptides to evaluate predictions: one where the excluded modification is encoded and one where it is not. Prediction performance for both test sets were then evaluated and compared. This comparison thus allows performance to be assessed on a modification that is not included in training, in terms of the improvement that DeepLC offers over a baseline of simply ignoring the presence of the modification.

Figures 4 and Supplementary Figure 6 show the prediction errors for each of the left-out modifications for training. In Figure 4, the boxplots show performance when a given modification was not present in the training set for the model, and is afterwards either not encoded (red boxplots; baseline) or encoded (blue boxplots) during the predictions. It should be noted that many modifications did not cause a substantial change in terms of predicted retention time, as was also observed in the original paper for this data set^25^ Examples of such modifications with limited impact are methyl, dimethyl, trimethyl, and deamidation. In contrast, the acyl modifications (including propionyl, succinyl, malonyl, crotonyl, and acetyl) show a clear performance increase when these modifications are encoded during the predictions. For instance, Supplementary Figure 6 shows that the MAE is improved by 700% (from 462 s to 66 s) for propionyl. These improvements are mainly due to the correct prediction by DeepLC of the shift in retention time caused by the modification, despite DeepLC never having encountered that specific modification before. Most importantly, besides a significantly decreased MAE, the correlation R also shows a substantial improvement. This is shown in Figure 4 through the substantially smaller variance for the blue box plots. For crotonyl, for instance, Supplementary Figure 6 presents an increase of R from 0.975 to 0.990 when encoding the modification in the test set. This means that the DeepLC models have not just learned the general shift in retention time caused by modifications, but also how this shift depends on the context of the modification in the peptide.

**Figure 4:**
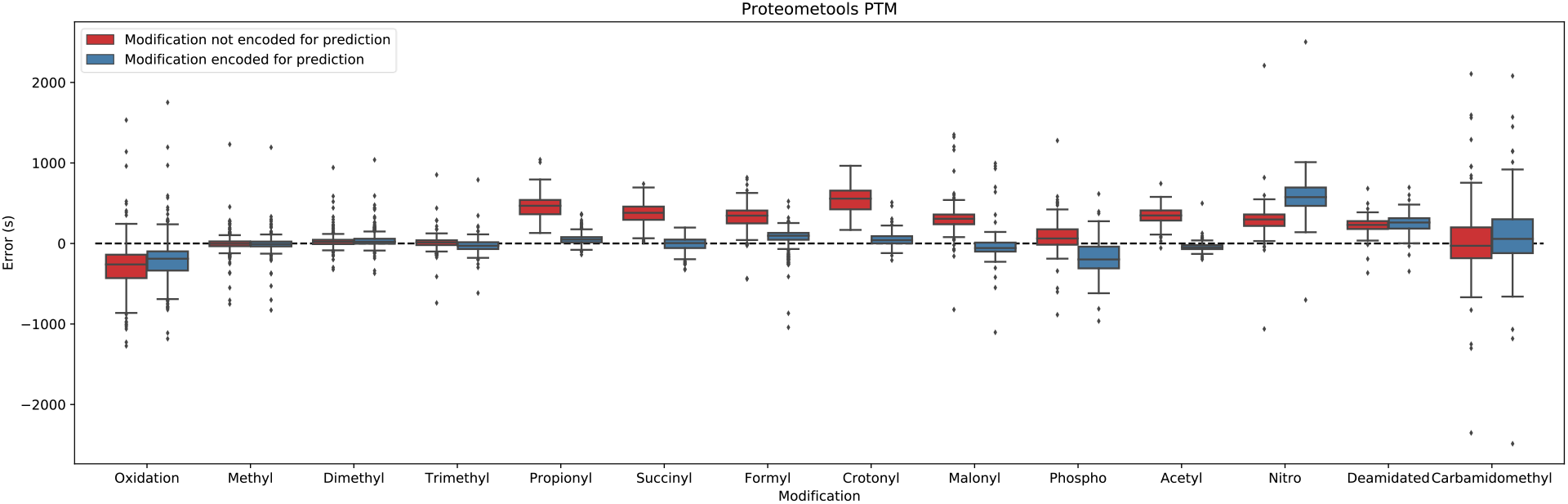
The modification that was excluded for training is shown on the horizontal axis, and the vertical axis shows the retention time error (experimental - predicted) when the modification was either not encoded (red) or encoded during the predictions (blue).

Only nitrotyrosine and phosphorylation modifications show lower performance when encoded, but these modifications can be classified as physicochemically very different from the other modifications. This inability of DeepLC to accurately predict retention times for modifications that are chemically very different from anything encountered the training set indicates that even DeepLC requires some relevant training data for a given class of modifications.

In the second evaluation procedure we used the larger *DIA HF* and the smaller *HeLa HF* data sets to train and optimize nineteen DeepLC models, where each model only sees peptides that do not contain a specific amino acid. The nomenclature is as above, in which model A is trained and optimized only on peptides that do not contain amino acid A. Next, each model is evaluated on all those peptides that do contain the excluded amino acid not seen during training. For this we again created two test sets from these remaining peptides: one where the excluded amino acid is encoded as the composition of glycine only and one with its actual composition.

We show that encoding an amino acid as itself instead of as glycine improves the MAE for most amino acids (Figure 5). DeepLC performs very well when modelling large hydrophobic residues as modified glycines, and slightly less well when modelling polar uncharged and negatively charged residues as modified glycines. Finally, for the positively charged amino acids only arginine shows an improvement, while lysine and histidine decrease in performance when encoding the amino acid. The poor performance for lysine can be explained by the difference with the amino acid with the closest atomic composition. For lysine the closest atomic compositions are arginine and leucine (or isoleucine), which are significantly less hydrophobic or more hydrophobic, respectively. This is similar to our observations for modifications, where the best performance is obtained for unseen modifications which can be more readily extrapolated from already seen modifications.

**Figure 5:**
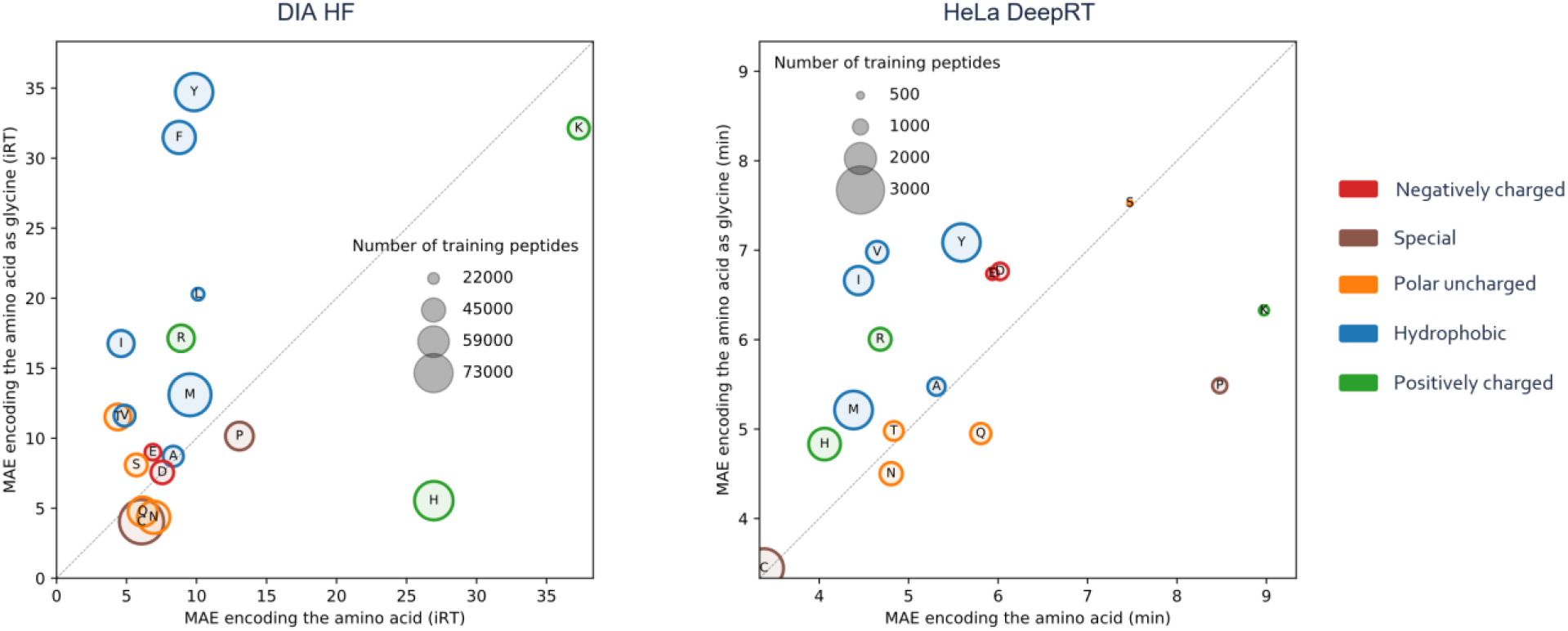
Each amino acid that was excluded for training is shown as a circle, where the size of the circle and color indicates the remaining training peptides and chemical property, respectively. The amino acid is either encoded as glycine (vertical axis) or as its own atomic composition (horizontal axis) and its position depicts the MAE for all amino acid containing peptides. This means that everything above the diagonal line is predicted with a higher accuracy when the amino acid is encoded as itself, while the reverse is true if it is below the diagonal line.

Between the larger and smaller data sets there is a consistent performance difference between the excluded amino acids. This consistency means that the observations made are likely to be independent of the specific data set the evaluation is run on.

This non-modified amino acid evaluation shows that performance is slightly worse in comparison to including the amino acid in the training set, with *DIA HF and HeLa DeepRT* having a MAE of 2.37 and 3.2 minutes, respectively. The MAE errors shown in Figure 5 are about 1.5 to 2.5 times higher.

It is important to note that this evaluation is not only hard because the trained model has never seen a given amino acid, it is also hard because peptides that are similar to each other are likely to all be excluded from training due to these peptides having a higher likelihood of also containing the removed amino acids. This can create biased training sets, especially for lysine and arginine as the majority of peptides are tryptic. Surprisingly, however, the model is still able to predict retention times very accurately for amino acids that were not used in training.

### Evaluation of open modification identifications

Predicted retention times have the potential to overcome the identification ambiguity issue^26,27^. Because of DeepLC’s unique capability to accurately predict retention times of (unseen) modifications, these predictions can be applied to open modification searches, where identification ambiguity is a key problem^21^. Indeed, open searches introduce considerable ambiguity through the very large number of possible modifications considered, which can be reduced through orthogonal measurements such as retention time.

DeepLC was applied on the results of an open modification search of CD8-positive T-cell data^28^ using the pFind^22^ search engine. Figure 6A shows observed retention time plotted against predicted retention time for the resulting peptide spectrum matches (PSMs). While the retention time is accurately predicted for PSMs with a Q-value lower than 0.01, much higher retention time errors are observed for PSMs with q-values higher than or equal to 0.01. Figure 6B shows that these PSMs do have a higher error, but the mode is still around zero. This indicates that we mostly expect false identifications to have their mode around zero with a large deviation from this mode.

**Figure 6:**
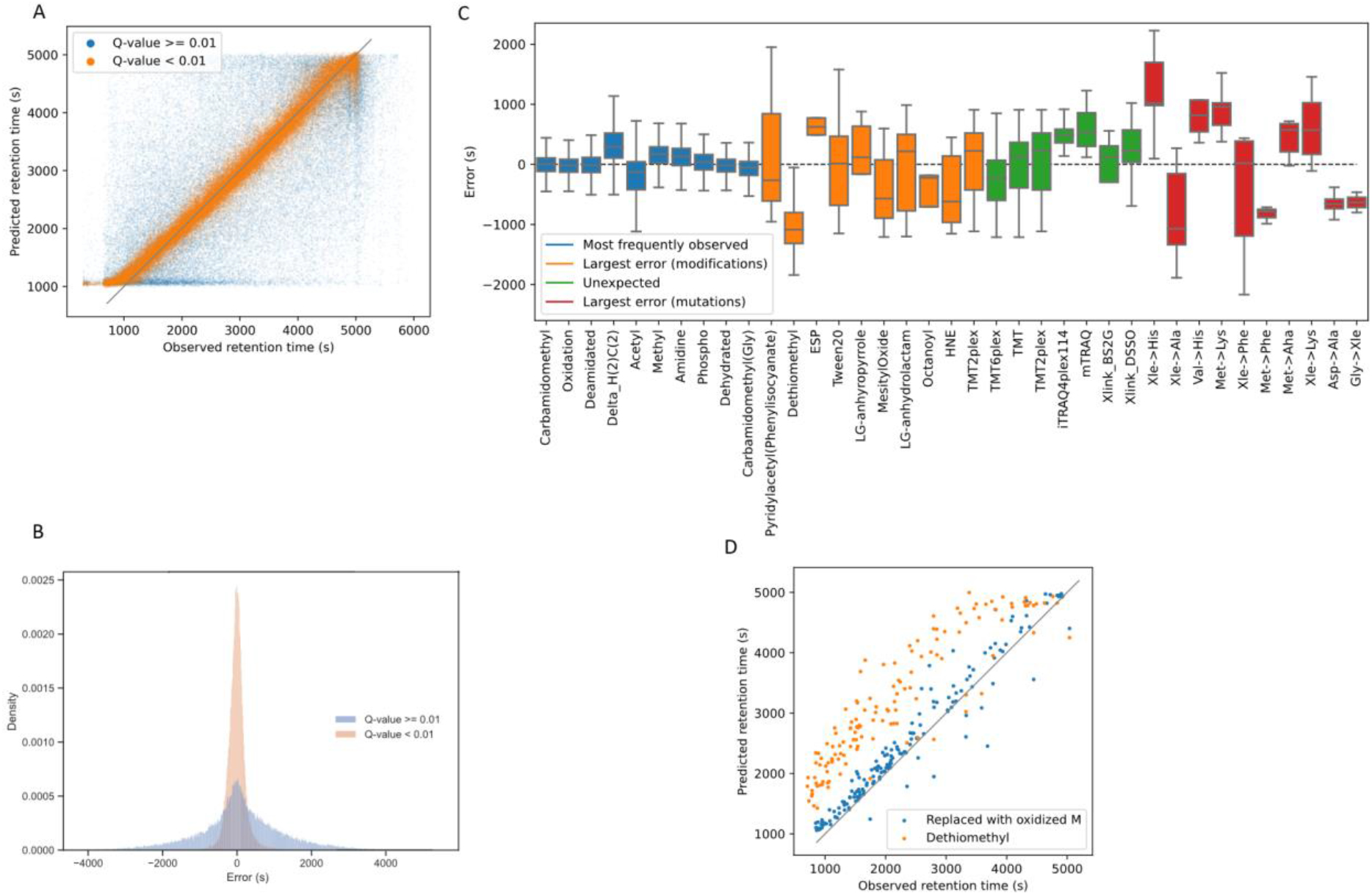
Predicted retention time analysis for open modification results of a CD8-positive T-cell data set. (A) Predicted and observed retention times split by Q-value. Panel (B) shows the error distribution split by Q-value and any points that are higher than 1.5 times the interquartile range plus the relevant quartile range are excluded from the plot. Panel (C) only shows PSMs with Q-values < 0.01. Colors indicate four different subsets of detected modifications. (1) PSMs carrying the top ten most abundant modifications (2) PSMs carrying the ten modifications with the largest absolute mean error (3) PSMs carrying modifications that are not expected to occur in the sample; and (4) PSMs carrying the top ten mutations with the largest absolute mean error. Panel (D) contains PSMs that were identified to contain a dethiomethyl modification and predictions for the same peptides where the dethiomethyl is replaced with an oxidation.

The error distribution of filtered PSMs (Q-value >= 0.01) is now compared to distributions of selected modifications to flag suspect modification groups. Boxplots of the error distributions for four subsets of modifications are shown in Figure 6C. The subset containing the ten most found modifications all show a low error spread that is within 5% of the maximum elution time (300 seconds) and is generally centered around zero. The subset of modifications with the largest errors are in the range of 25% of the maximum elution time (1500 seconds) and are mostly centered around zero seconds. A notable exception is dethiomethyl with a shifted median retention time error of 1000 seconds. This shift can be explained by in-source fragmentation, which causes oxidized methionine to lose its side chain^29^. If in-source fragmentation is the cause than the observed retention times are expected to be based on the oxidized methionine equivalent. To verify whether this is the case, PSMs with dethiomethyl were replaced with their oxidized methionine precursors. Figure 6D shows that replacing detiomethyl with oxidized methionine reduces the predicted retention time error to around the same level as expected for oxidized methionine peptides in Figure 6C.

The subset of modifications with large errors shows very similar patterns to the next subset, which contains modifications that are not expected to occur in the sample, as these are experimentally induced and thus should not be found in the untreated biological sample under analysis here. In effect, the similarity between these two subsets of modifications indicates that modifications with large retention time errors according to DeepLC can be flagged as highly suspect.

Finally, the last subset singles out presumed detection of mutations, which show similar or worse error distributions than the previous two subsets. The large error hints at an incorrect interpretation of the observed mass shift. For these incorrect interpretations one could expect bimodal distributions that contain a combination of right and wrongly assigned mass shifts, but the error distributions are mostly unimodal. The observation that presumed mutations are among the most problematic corresponds to the known non-trivial nature of reliably identifying such sequence changes^26,30^.

These results thus demonstrate that the unique capabilities of DeepLC allow it to be used as an orthogonal measure to flag suspect identifications in open modification searches.

## Discussion & Conclusion

Our evaluation shows that DeepLC performs similarly to current state-of-the-art models for unmodified peptides. DeepLC performance furthermore increases for larger data sets, where models trained on larger data sets can provide accurate predictions for smaller data sets. More importantly, DeepLC can accurately predict the retention time of modified peptides, even for modifications that were not included in the training set. This ability to predict for unseen modifications was evaluated with a two-pronged evaluation strategy using both unmodified peptides as well as synthetic, modified peptides. For both evaluations encoding modifications for prediction improves performance, while performance is reduced only for specific modifications that are very different from any other structure in the data set. Finally, the potential of this unique capability of DeepLC is illustrated through its ability to flag suspect PSMs in an open modification search. Crucially, DeepLC showed much larger prediction errors for PSMs that carried modifications that are certain to be absent from the sample.

Future development of models that can predict the retention time for unseen modifications could focus on structural aspects of modifications. DeepLC is currently limited in differentiating between isomeric structures that are physicochemically different. Indeed, the observation that structure, not only atomic composition, leads to the physicochemical properties of molecules has already been observed for small molecules. Here, the decision was made to work with atomic composition, because of the ready availability of the composition in databases like Unimod, and greater ease of integration when compared to more complex structural descriptors.

DeepLC enables the field to generate predictions for a wide landscape of modification. In order to improve the availability to researchers and their use-cases, DeepLC is made freely available online and has a user-friendly GUI. Furthermore, the tool is available in code repositories that enable easy incorporation in workflows and pipelines for automatic predictions.

## Methods

### Architecture

DeepLC uses a convolutional deep learning architecture with four different paths for a given encoded peptide. The same peptide acts as the input for the four paths, which have multiple separated layers, as shown in Figure 7. Three of the initial paths use a combination of convolutional^31^ and max pooling layers^32^. The remaining path, which propagates global features, consists of densely connected layers. The results of all initial four paths are flattened and concatenated to provide an input for the final combined path which consists of six connected dense layers. A detailed visualization of the architecture is available in Supplementary Figure 7.

**Figure 7:**
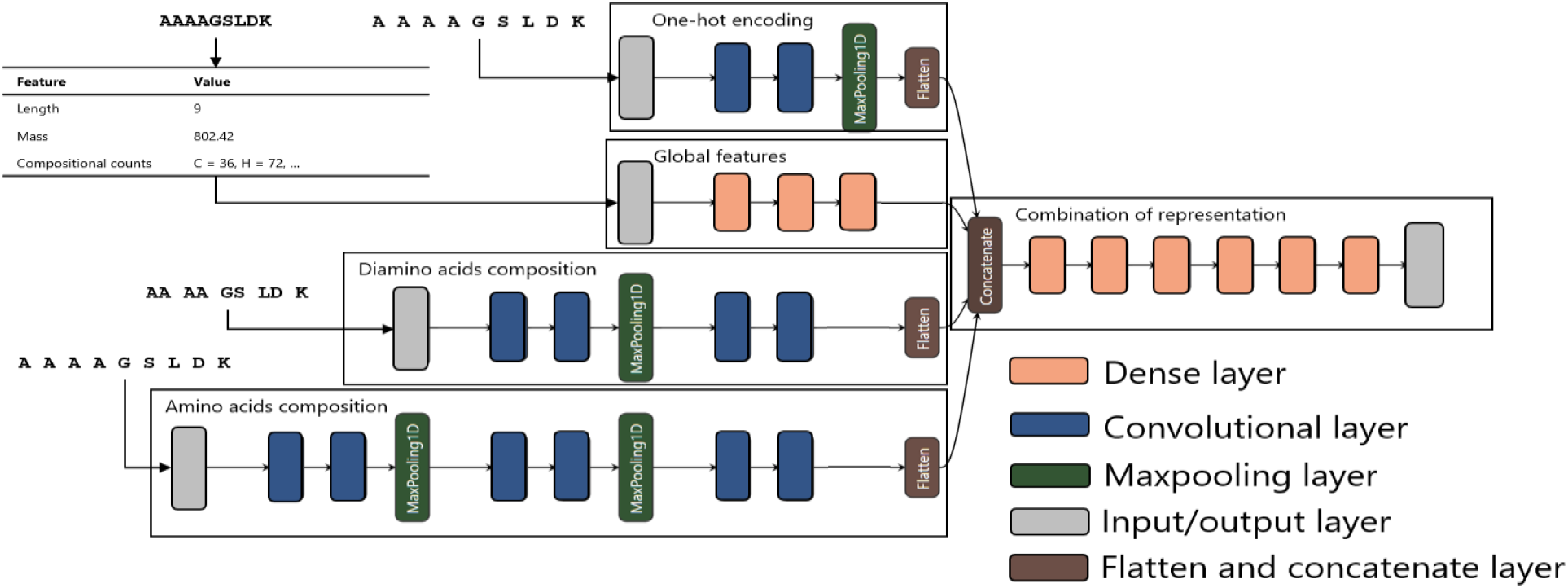
Visualization of DeepLC’s convolutional architecture with the four individual paths named: *One-hot encoding, Global features, Diamino acids composition*, and *Amino acids composition*. These individual paths are concatenated in the *Combination of representations* path.

The input matrix for the *Amino acids composition* path has a dimension of 60 for the peptide sequence by 7 for the atom counts (C, H, N, O, P, and S). Not every peptide is 60 amino acids long, thus “X”-characters without atomic composition are padded to reach 60 amino acids. This implies that encoding modified amino acids becomes straightforward, as computing their atomic composition is trivial. Note that for modified amino acids the atomic composition of the modification is added to the atomic composition of the unmodified residue. This encoding allows the model to learn patterns that generalize to unseen modifications.

The *Diamino acid* path was added to further improve the generalization capability of the model. In this layer the peptide is divided into diamino acids without overlap. This improves the generalization capability, as the input values for each position are more thoroughly represented. Otherwise there would only be 20 unmodified amino acid representations, combined with a limited amount of modifications. Besides interpreting the amino acids in pairs, the *Diamino acid* path uses the same logic as the *Amino acids composition* path, leading to an input matrix of 30 paired positions by 6 atoms.

Encoding amino acids and their modifications by strictly using the atomic composition does, however, not allow for comprehensively capturing all molecular information. Indeed, the structure of isomers can play an important role in the physicochemical properties of amino acids, as is exemplified by structural isomers isoleucine and leucine^33^. This is the reason that one-hot encoding of unmodified amino acids was still used in DeepLC as an input for the *One-hot encoding* path. However, to reduce the impact of this layer, the number of filters for this path were limited to a mere 2. The dimensions of this input matrix is 60 positions by 20 amino acids.

In addition to all paths that encode position specific information, the *Global features* path takes global information of the peptide into account. These global features include the length and total atomic composition of the peptide. The dimension of this input vector is 7.

Three versions of the model were trained, solely differing in kernel size (of 2, 4, and 8) for the *Amino acids composition* path. These three models were combined in an ensemble by averaging their predictions.

Finally, the other hyperparameters of each layer in DeepLC are consistent for all versions with different kernel sizes. All layers, except the output layer and the *One-hot encoding* path, use L1 regularization with *α* = 2.5*e* – 7 and a leaky ReLU^34^ with a maximum activation value of 20. The *One-hot encoding* path uses the tanh activation function, as within this path we are only interested in the ability to separate unmodified amino acid isomers.

### Data sets & evaluation

To evaluate the generalization performance of DeepLC, we selected 20 data sets from a wide variety of organisms and experimental setups (Supplementary Table 2). We further selected three data sets (*SWATH Library*^35^, *HeLa HF*^36^, and *DIA HF*^37^) for detailed result reporting, with the results for the other 17 data sets described in the supplementary information. The data sets *SWATH Library* and *DIA HF* were selected based on their previous use by Ma *et al*. for DeepRT^11^ and by Guan *et al*.^9^, respectively. A third data set, *HeLa HF* was selected because of its use of short (compared to other used data sets) gradients of 15 minutes and the large number of training peptides.

The variety in experimental setups and protocols means that the acquired and predicted retention times need to be calibrated. The *ProteomeTools library*^38^, *SWATH Library*, and *DIA HF* data sets were normalized to the iRT peptides^39,40^. DeepLC itself supports linear calibration which is further explained in the online DeepLC documentation.

The data sets marked *Custom workflow* in Supplementary Table 2 were processed as follows. Raw mass spectrometry files were downloaded from PRIDE Archive^41^ and converted to MGF format with the ThermoRawFileParser^42^. These were then searched using the MS-GF+ search engine^43^ with a concatenated target-decoy sequence database containing the respective species’ UniProtKB proteome and the common Repository of Adventitious Proteins (cRAP: https://www.thegpm.org/crap/). The MS-GF+ search results were post-processed with Percolator^44^ to a false discovery rate of 0.01. Retention times were parsed from the MGF files for all confidently identified peptides. Within each LC-MS run, the median retention time for each peptidoform (peptide - modifications combination) was calculated. All median retention times were then linearly calibrated across all LC-MS runs for each data set, using the shared peptidoforms as anchor points. Finally, the median calibrated retention time was calculated for each peptidoform across all runs for each data set. These median calibrated retention times were then used to train, validate and test DeepLC. The steps comprising this calibration are available in a Snakemake workflow^45^.

The data sets marked *Custom workflow ProteomeTools* in Supplementary Table 2 were processed as follows. MaxQuant^46^ identification files were filtered on posterior error probabilities < 0.01 and scores > 90. The retention times were calibrated with the peptides in Supplementary Table 3. Within a run, the median retention time per peptidoform was used for further analysis. Then, across runs the median retention time per peptidoform was taken for the final retention time. The data sets marked *Custom workflow ProteomeTools PTM*^25^ in Supplementary Table 2 were processed in the same way as the data sets *Custom workflow ProteomeTools*, with the only exception that the retention times were calibrated with the peptides in Supplementary Table 4. Each data set was split into a test set (10%), validation set (5%), and training set (85%). The validation set is used for model selection only, while all performance results presented here were computed from the test set. Prediction performance is measured using three commonly used metrics: Mean Absolute Error (MAE), Pearson correlation, and Δ*t*_95%_. The latter describes the error for a retention time window that contains 95% of the peptides in the error distribution. To make the MAE and Δ*t*_95%_ comparable between experiments, we divided them by the retention time of the last detected peptide in the respective data set. These metrics are further referred to as relative MAE and relative Δ*t*_95%_.

### Open modification search

Results from an open modification search by pFind^26^ are used to evaluate the ability of DeepLC to identify suspect identifications. Open modification searches are particularly prone to falsely identify modified peptides due to the larger search space and resulting identification ambiguity^46^. The search was performed on a data set on CD8-positive T cells by Kim *et al*.^28^, obtained from the PRIDE repository^47^ with identifier PXD000561. The search was run with pFind version 3.1.5. Search parameters were set to: 20ppm precursor mass error tolerance, peptide length limit from 7 to 30 amino acids, modifications were limited to mass deltas from -150 to 500 Dalton, a maximum of two miss-cleavages were allowed, and oxidation of M and carbamidomethylation of C were set as variable modifications.

### Software and scripts

The following Python libraries were used in DeepLC: Pandas^48^, TensorFlow^49^, Pyteomics^50^, SciPy^51^, and Numpy^52^. The code used to prepare the data sets, calibrate retention times, generate DeepLC models, make predictions, and generate the figures is available on Zenodo: https://zenodo.org/record/4542884.

For the DeepLC tool (Supplementary Figure 8) itself, see Availability.

## Availability

DeepLC is available for download from the following repositories and package indexes:

– Graphical user interface: https://github.com/compomics/DeepLC/releases/latest
– Python package: https://pypi.org/project/deeplc/
– Bioconda package: https://bioconda.github.io/recipes/deeplc/README.html
– Biocontainers docker image: https://quay.io/repository/biocontainers/deeplc
– Source code: https://github.com/compomics/DeepLC

## Acknowledgements

R.B. acknowledges funding from the Marie Sklodowska-Curie EU Framework for Research and Innovation Horizon 2020 MASSTRPLAN [675132] and Vlaams Agentschap Innoveren en Ondernemen under project number HBC.2020.2205; R.G. acknowledges funding from the Research Foundation Flanders (FWO) [1S50918N]; S.D. and L.M. acknowledge funding from the European Union’s Horizon 2020 Programme (H2020-INFRAIA-2018-1) [823839]; N.H. and L.M. acknowledge funding from the Research Foundation Flanders (FWO) [G042518N].

## Contributions

R.B., R.G., and S.D. conceived the study. R.B., R.G., L.M. and S.D. designed the experiments, analyzed the results, and wrote the manuscript. R.G. made the tool available in python package repositories. N.H. and R.B. build the graphical user interface.

## Supplemental information

**Supplemental Figure 1:**
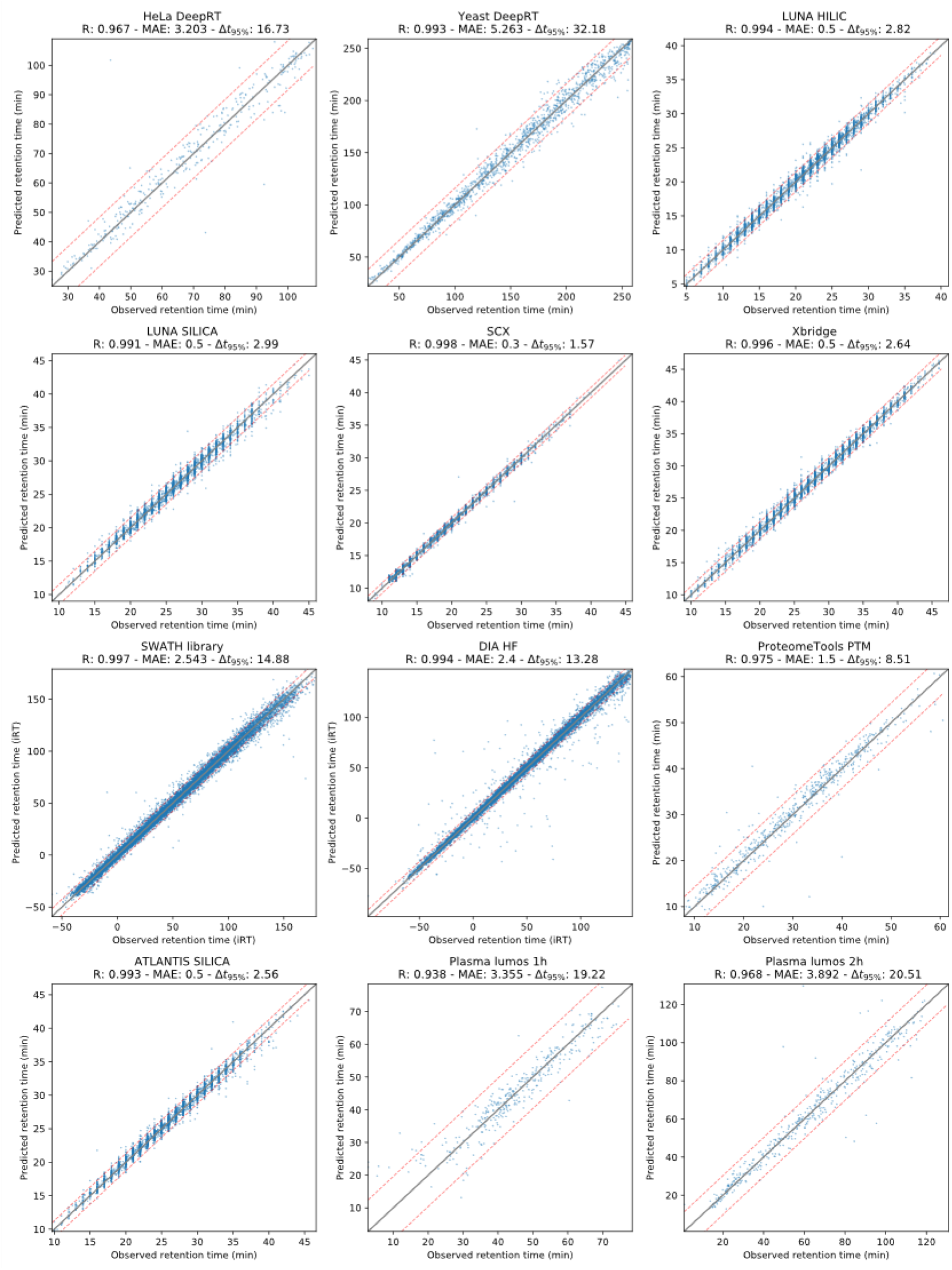
Scatter plots for twelve data sets on the test data for each respective data set.

**Supplemental Figure 2:**
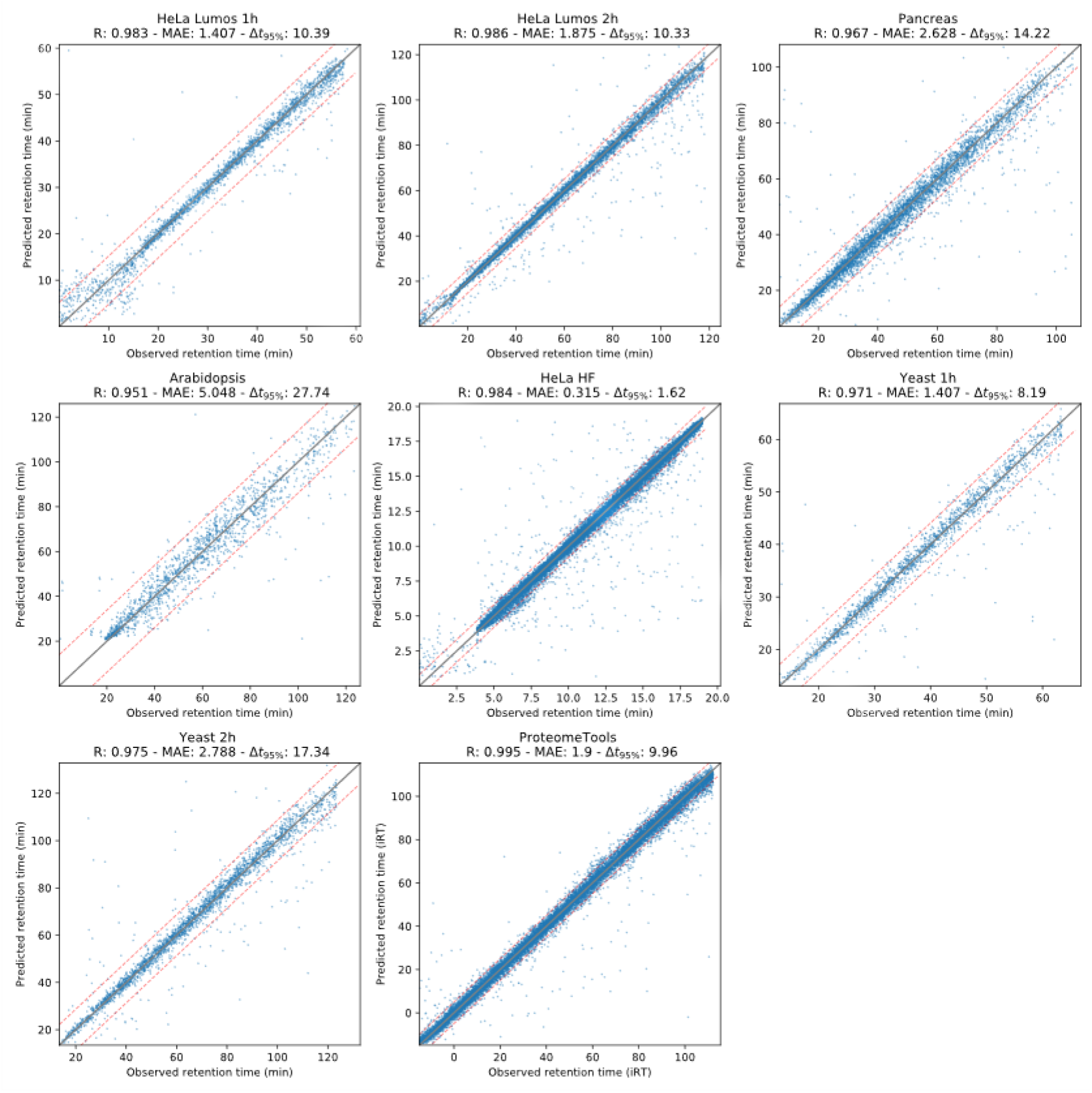
Scatter plots for eight data sets on the test data for each respective data set.

**Supplemental Figure 3:**
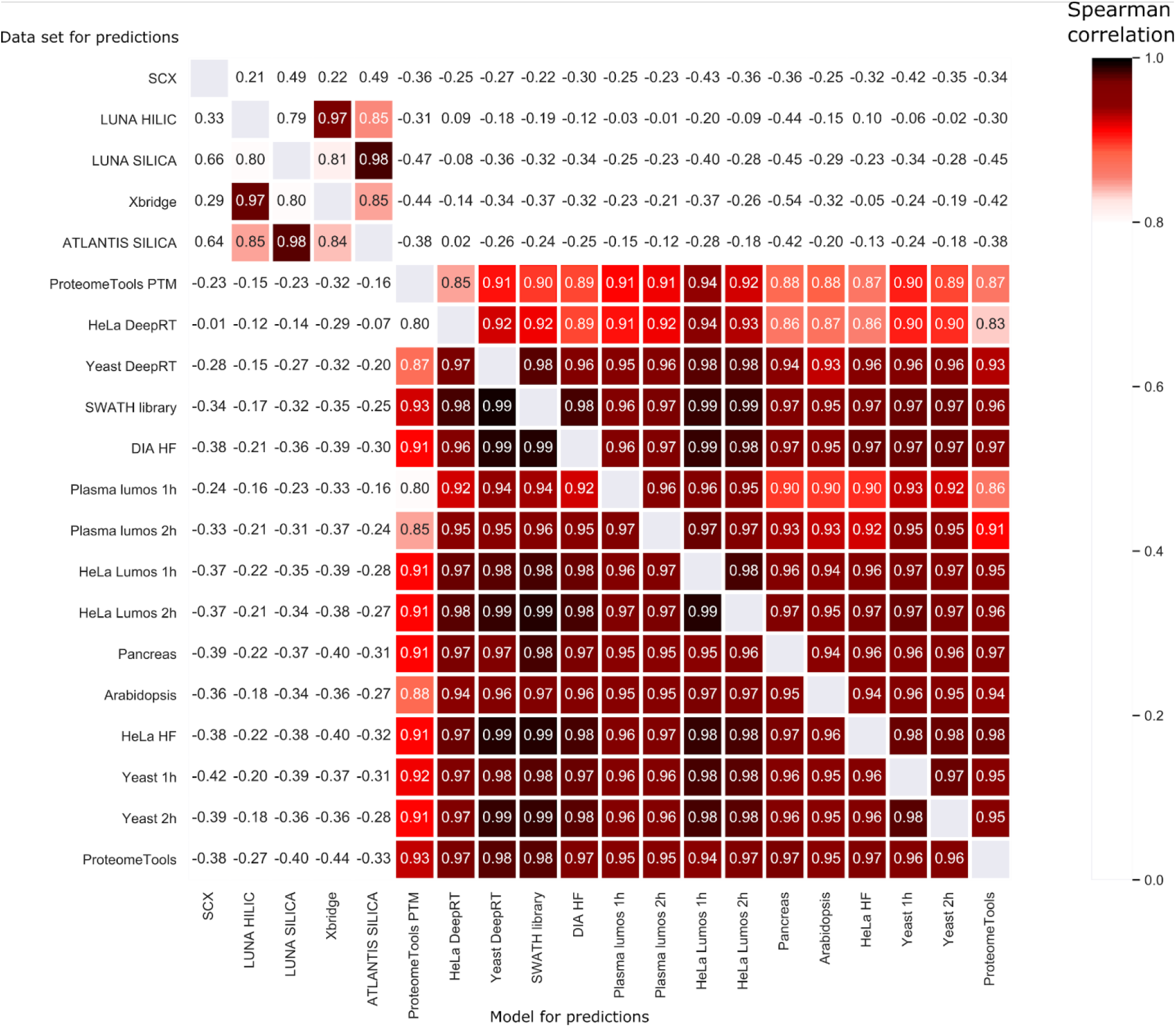
Performance in terms of the Spearman correlation for each model applied to the other 19 data sets.

**Supplemental Figure 4:**
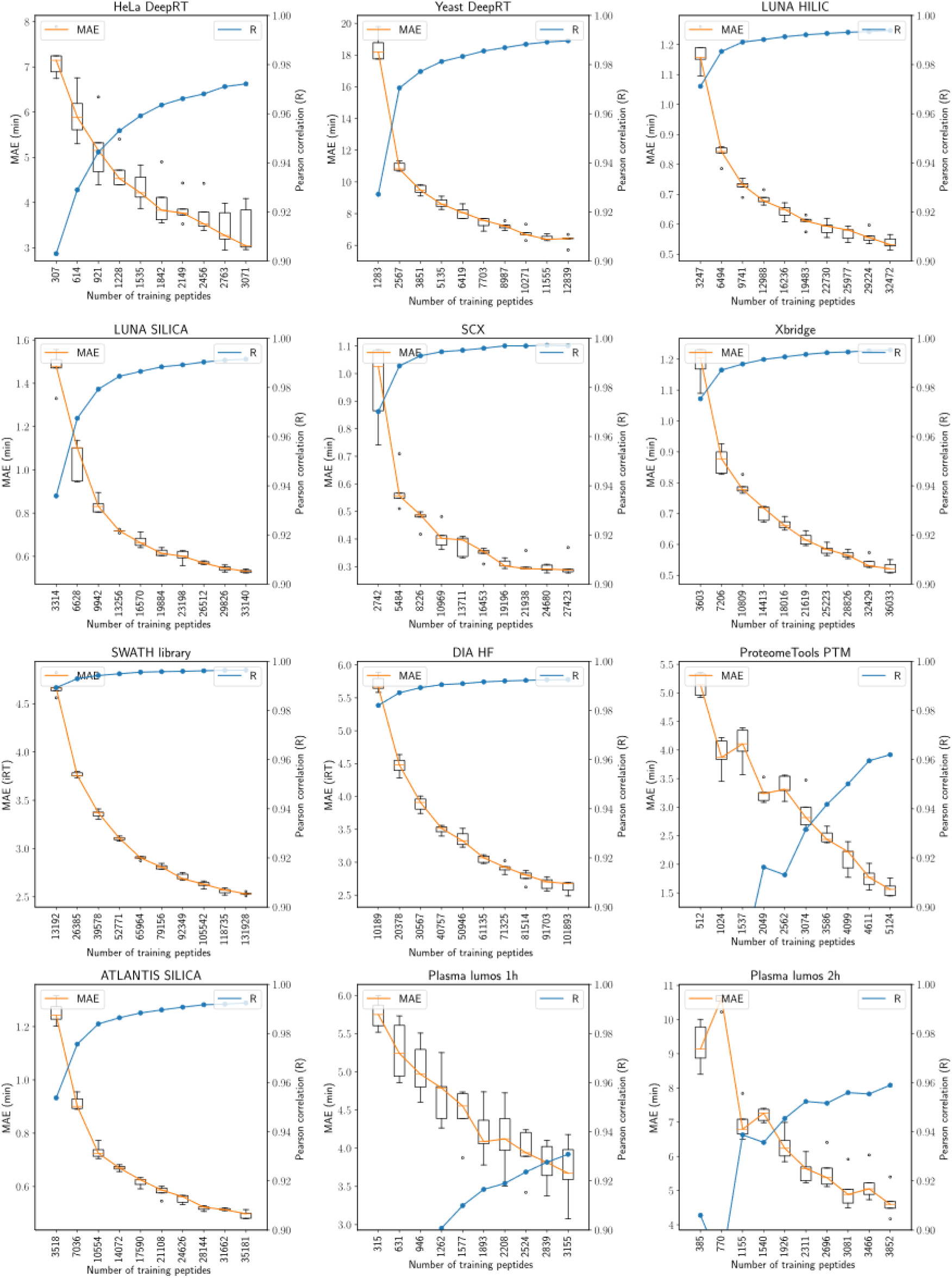
Learning curves for twelve data sets with the MAE and correlation between observed and predicted retention times.

**Supplemental Figure 5:**
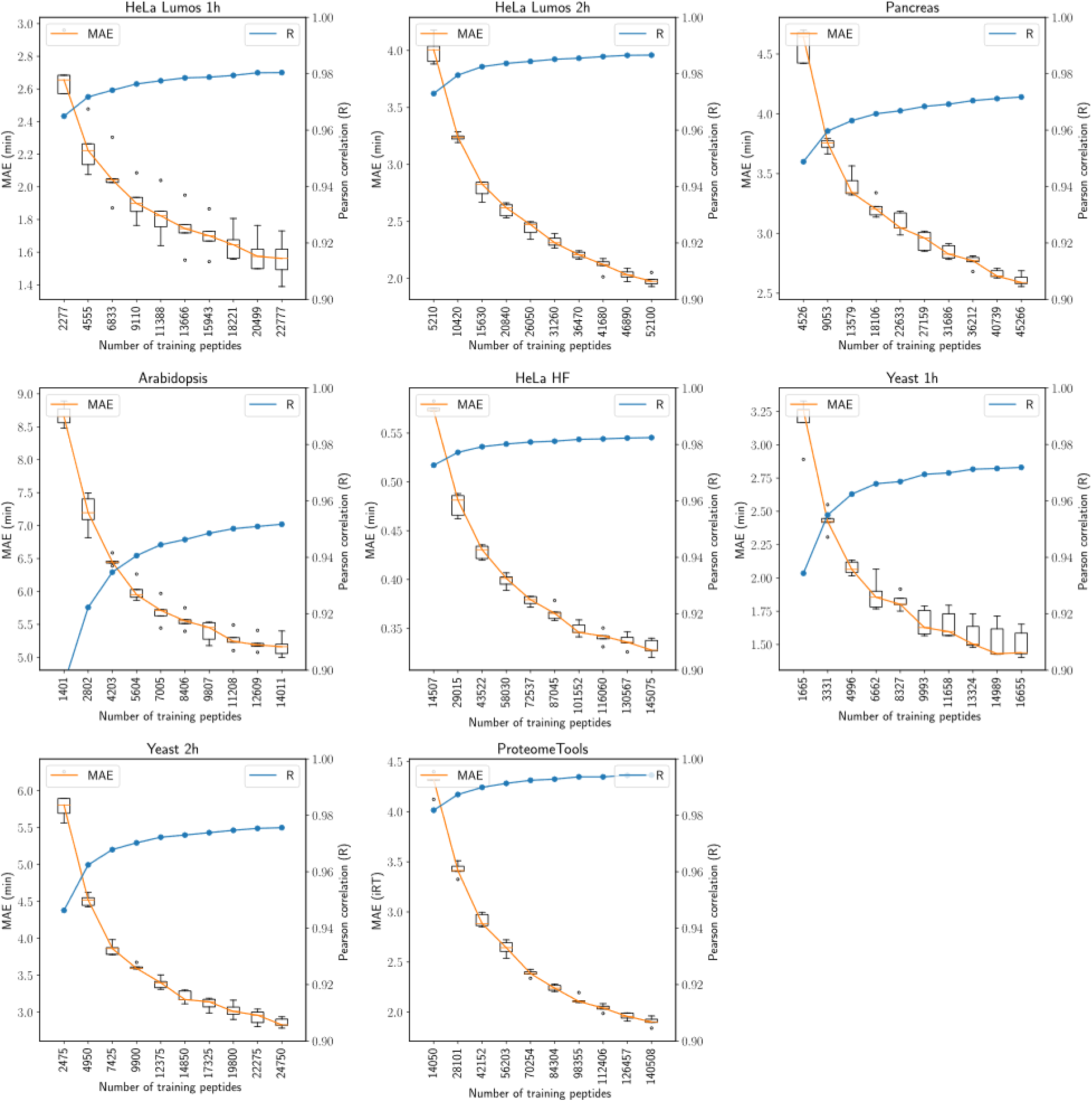
Learning curves for eight data sets with the MAE and correlation between observed and predicted retention times.

**Supplemental Table 1:**
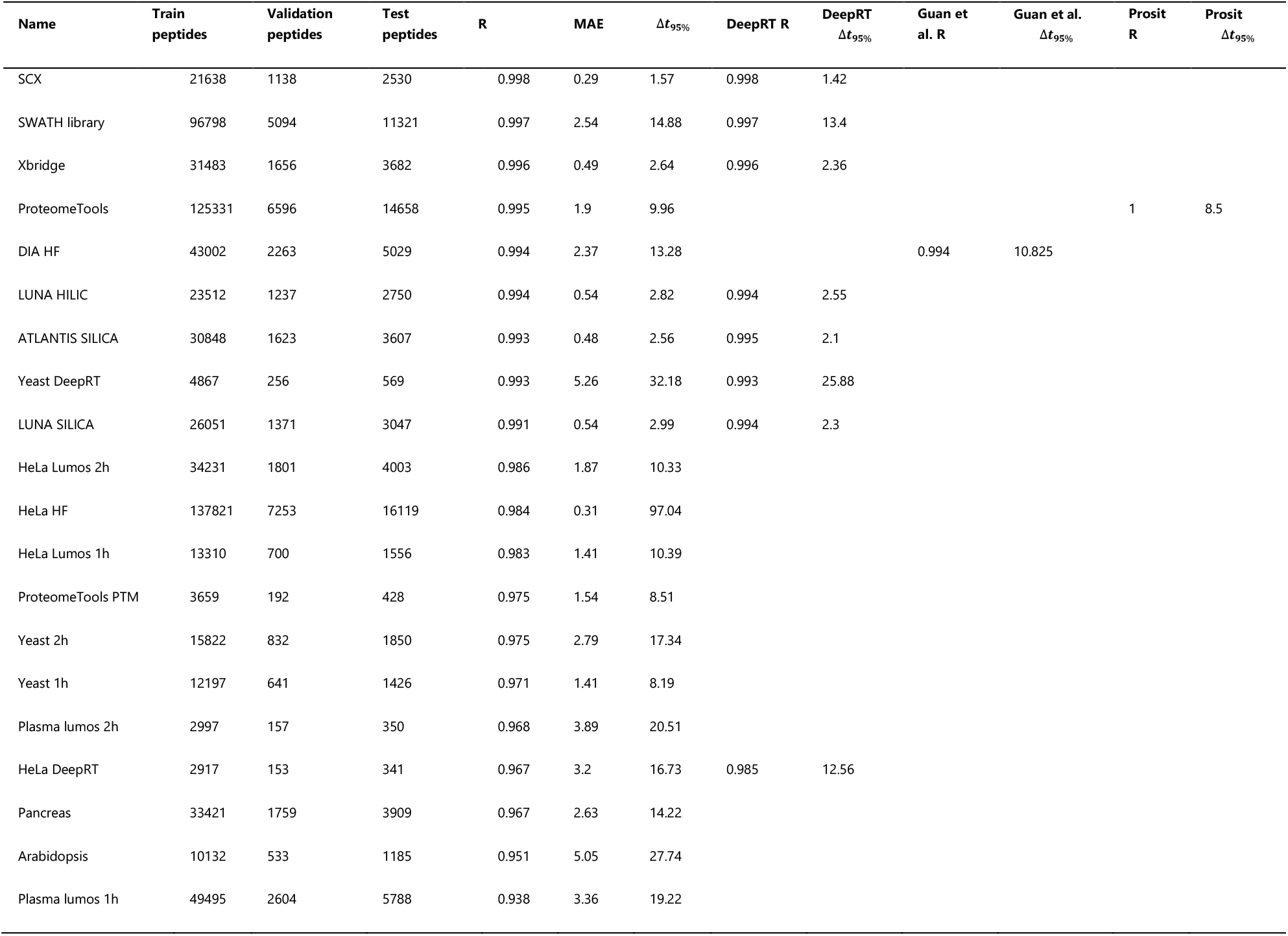
Overview of the 20 MS2 data sets used in this research. The first three columns report test performance for DeepLC. The performance for existing models is listed if available and as reported in the original manuscript (except for the Prosit model where the Δ*t*_95%_ was recalculated based on a digitized version of Figure 1c in the original manuscript). For Guan et al. the Δ*t*_95%_ was recalculated by taking the Δ*t*_95%_ of the absolute error and multiplying this value by two. This value was corrected to be consistent with DeepRT and DeepLC.

| Name | Train peptides | Validation peptides | Test peptides | R | MAE | $\Delta t_{95\%}$ | DeepRT R | DeepRT $\Delta t_{95\%}$ | Guan et al. R | Guan et al. $\Delta t_{95\%}$ | Prosit R | Prosit $\Delta t_{95\%}$ |
| --- | --- | --- | --- | --- | --- | --- | --- | --- | --- | --- | --- | --- |
| SCX | 21638 | 1138 | 2530 | 0.998 | 0.29 | 1.57 | 0.998 | 1.42 |  |  |  |  |
| SWATH library | 96798 | 5094 | 11321 | 0.997 | 2.54 | 14.88 | 0.997 | 13.4 |  |  |  |  |
| Xbridge | 31483 | 1656 | 3682 | 0.996 | 0.49 | 2.64 | 0.996 | 2.36 |  |  |  |  |
| ProteomeTools | 125331 | 6596 | 14658 | 0.995 | 1.9 | 9.96 |  |  |  |  | 1 | 8.5 |
| DIA HF | 43002 | 2263 | 5029 | 0.994 | 2.37 | 13.28 |  |  | 0.994 | 10.825 |  |  |
| LUNA HILIC | 23512 | 1237 | 2750 | 0.994 | 0.54 | 2.82 | 0.994 | 2.55 |  |  |  |  |
| ATLANTIS SILICA | 30848 | 1623 | 3607 | 0.993 | 0.48 | 2.56 | 0.995 | 2.1 |  |  |  |  |
| Yeast DeepRT | 4867 | 256 | 569 | 0.993 | 5.26 | 32.18 | 0.993 | 25.88 |  |  |  |  |
| LUNA SILICA | 26051 | 1371 | 3047 | 0.991 | 0.54 | 2.99 | 0.994 | 2.3 |  |  |  |  |
| HeLa Lumos 2h | 34231 | 1801 | 4003 | 0.986 | 1.87 | 10.33 |  |  |  |  |  |  |
| HeLa HF | 137821 | 7253 | 16119 | 0.984 | 0.31 | 97.04 |  |  |  |  |  |  |
| HeLa Lumos 1h | 13310 | 700 | 1556 | 0.983 | 1.41 | 10.39 |  |  |  |  |  |  |
| ProteomeTools PTM | 3659 | 192 | 428 | 0.975 | 1.54 | 8.51 |  |  |  |  |  |  |
| Yeast 2h | 15822 | 832 | 1850 | 0.975 | 2.79 | 17.34 |  |  |  |  |  |  |
| Yeast 1h | 12197 | 641 | 1426 | 0.971 | 1.41 | 8.19 |  |  |  |  |  |  |
| Plasma lumos 2h | 2997 | 157 | 350 | 0.968 | 3.89 | 20.51 |  |  |  |  |  |  |
| HeLa DeepRT | 2917 | 153 | 341 | 0.967 | 3.2 | 16.73 | 0.985 | 12.56 |  |  |  |  |
| Pancreas | 33421 | 1759 | 3909 | 0.967 | 2.63 | 14.22 |  |  |  |  |  |  |
| Arabidopsis | 10132 | 533 | 1185 | 0.951 | 5.05 | 27.74 |  |  |  |  |  |  |
| Plasma lumos 1h | 49495 | 2604 | 5788 | 0.938 | 3.36 | 19.22 |  |  |  |  |  |  |

**Supplemental Figure 6:**
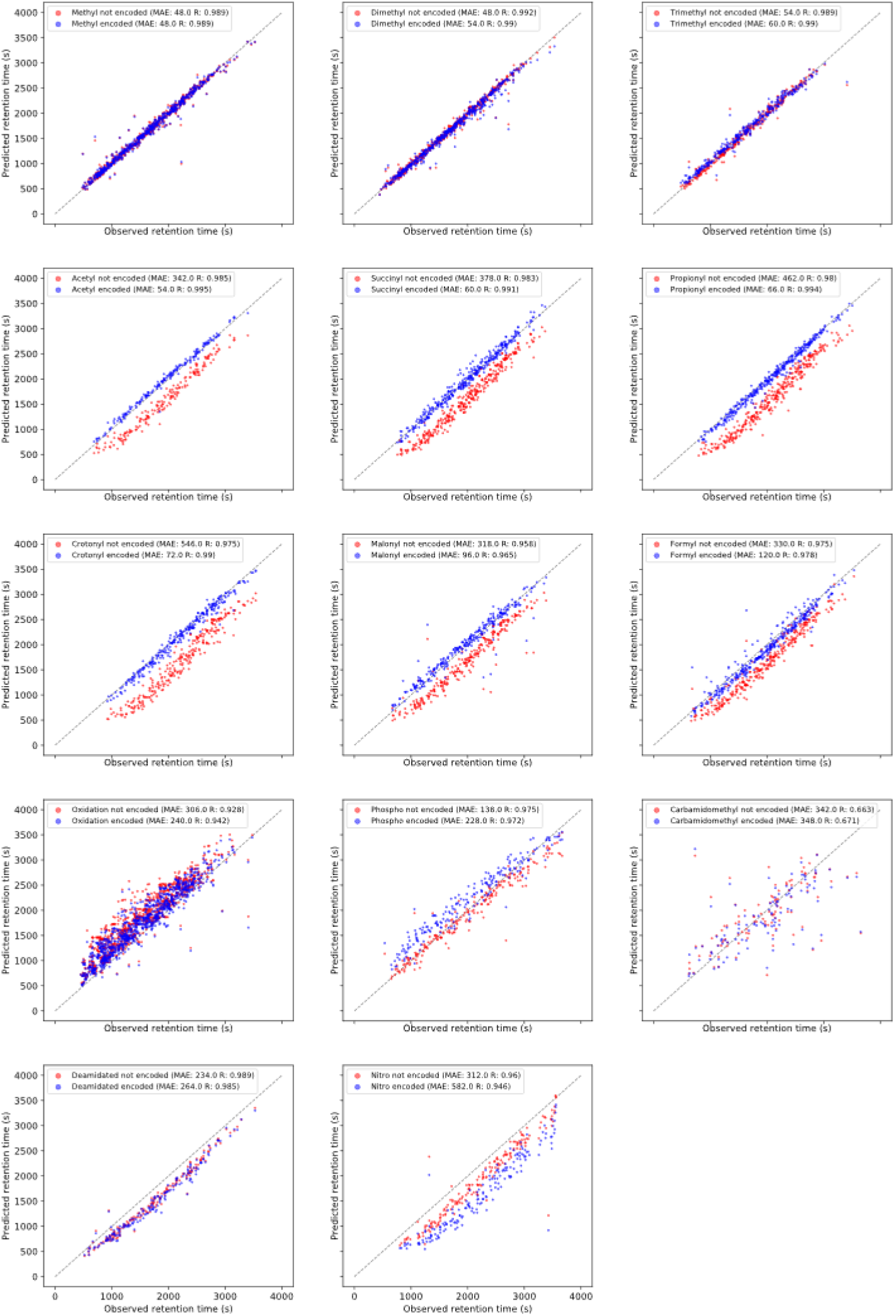
Each panel shows the observed retention time against predicted retention time for models that were not trained for the specified modification. Dots show the retention times when modifications are either not encoded (red) or encoded (blue) by DeepLC.

**Supplemental Figure 7:**
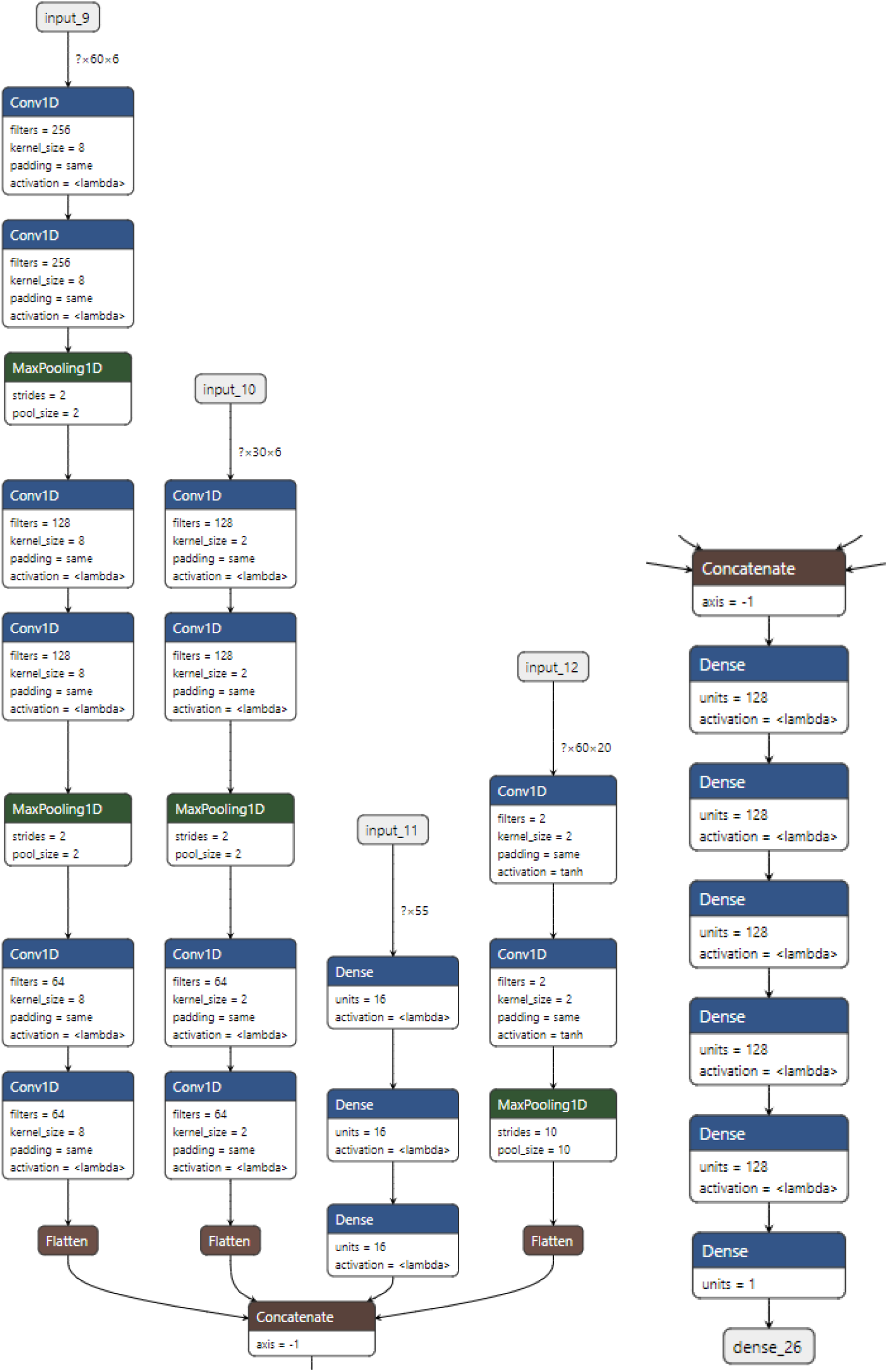
Detailed view of the architecture of DeepLC.

**Supplemental Figure 8:**
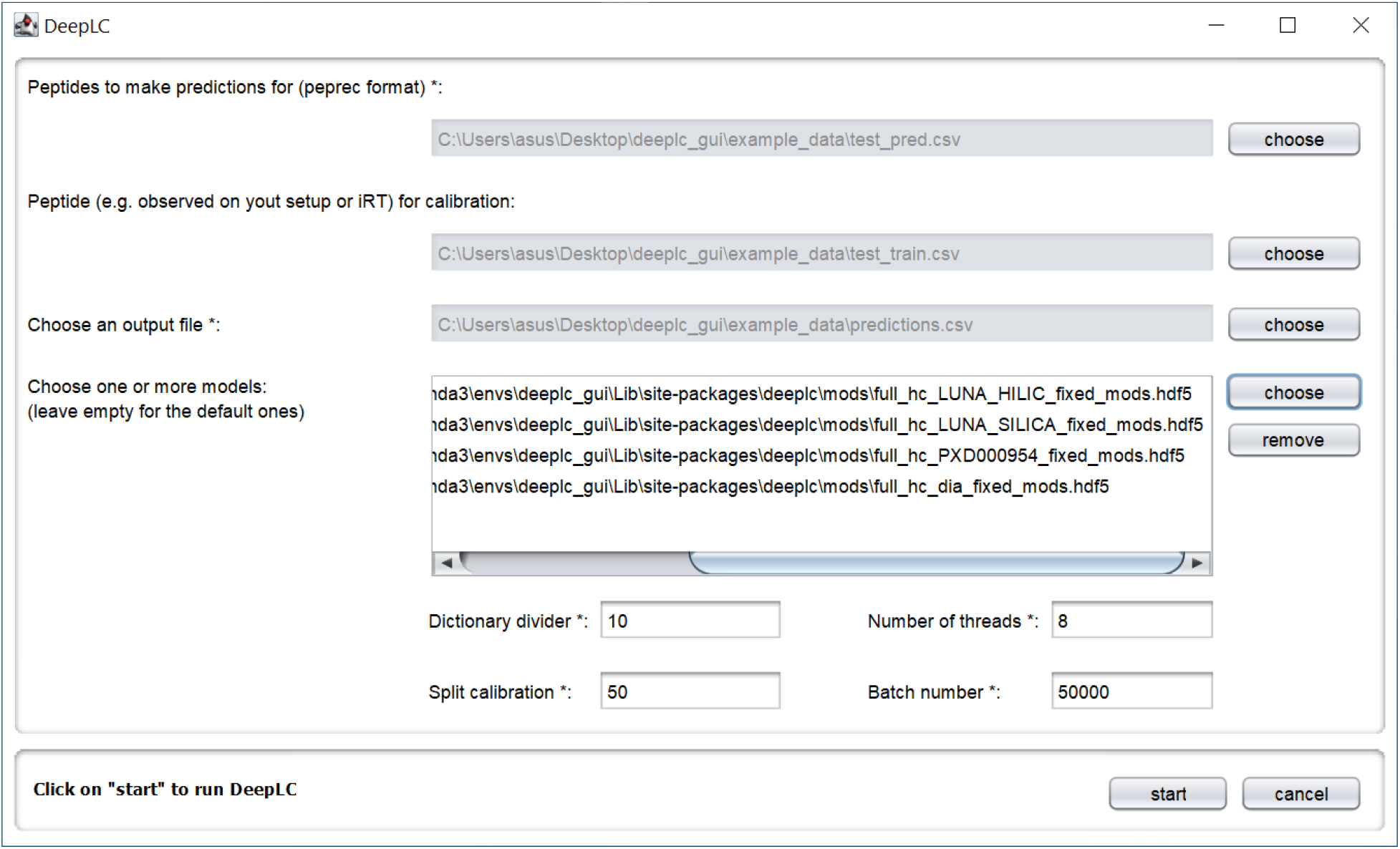
Java-based graphical user interface of DeepLC, which allows an easy to use interface to make predictions, with the option to calibrate for the user’s experimental setup.

**Supplemental Figure 9:**
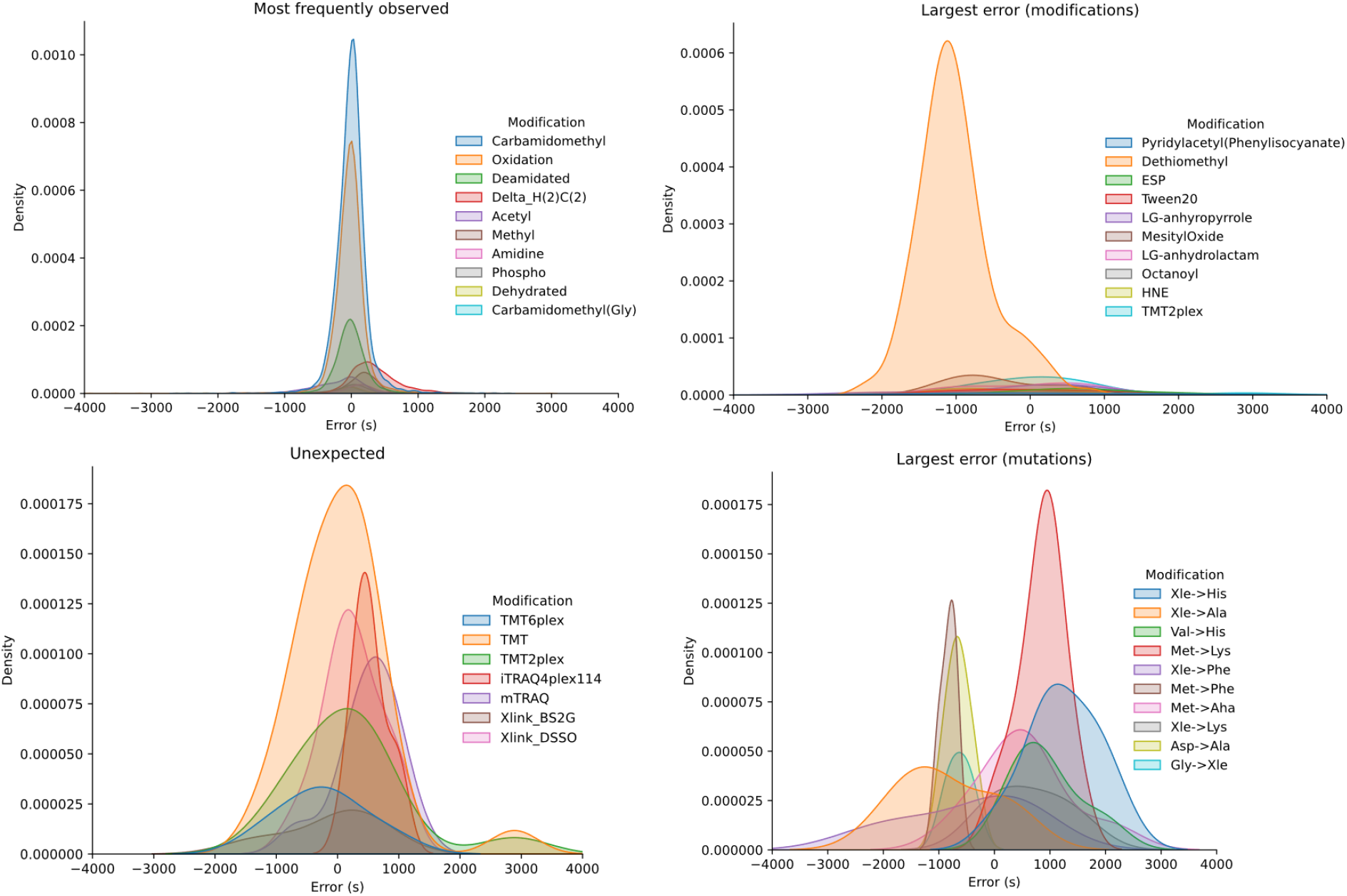
Error distribution for four different sets of modifications. (1) PSMs carrying the top ten most abundant modifications (2) PSMs carrying the ten modifications with the largest absolute mean error (3) PSMs carrying modifications that are not expected to occur in the sample; and (4) PSMs carrying the top ten mutations with the largest absolute mean error.

**Supplemental Table 2:** Data sets used to train and evaluate DeepLC

| Name | Data origin for fitting | Training peptides | Validation peptides | Test peptides | Repository identifier | Column type |
| --- | --- | --- | --- | --- | --- | --- |
| HeLa hf<sup>30</sup> | Custom workflow | 137821 | 7253 | 16119 | PXD006932 | RP |
| ProteomeTools<sup>32</sup> | Custom workflow<br>ProteomeTools | 125331 | 6596 | 14658 | PXD010595; PXD004732 | RP |
| SWATH library<sup>29</sup> | DeepRT<sup>11</sup> | 96798 | 5094 | 11321 | PXD000954 | RP |
| Plasma lumos 1h<sup>47</sup> | Custom workflow | 49495 | 2604 | 5788 | PXD013477 | RP |
| DIA HF<sup>31</sup> | Guan et al.<sup>9</sup> | 43002 | 2263 | 5029 | PXD005573 | RP |
| HeLa lumos 2h<sup>47</sup> | Custom workflow | 34231 | 1801 | 4003 | PXD013477 | RP |
| Pancreas<sup>48</sup> | Custom workflow | 33421 | 1759 | 3909 | PXD010154 | RP |
| Xbridge<sup>49</sup> | DeepRT<sup>11</sup> | 31483 | 1656 | 3682 |  | HILIC |
| ATLANTIS SILICA<sup>49</sup> | DeepRT<sup>11</sup> | 30848 | 1623 | 3607 |  | HILIC |
| LUNA SILICA<sup>49</sup> | DeepRT<sup>11</sup> | 26051 | 1371 | 3047 |  | HILIC |
| LUNA HILIC<sup>49</sup> | DeepRT<sup>11</sup> | 23512 | 1237 | 2750 |  | HILIC |
| SCX<sup>49</sup> | DeepRT<sup>11</sup> | 21638 | 1138 | 2530 |  | SCX |
| Yeast 2h<sup>50</sup> | Custom workflow | 15822 | 832 | 1850 | PXD003472 | RP |
| HeLa lumos 1h<sup>47</sup> | Custom workflow | 13310 | 700 | 1556 | PXD013477 | RP |
| Yeast 1h<sup>50</sup> | Custom workflow | 12197 | 641 | 1426 | PXD003472 | RP |
| Arabidopsis<sup>51</sup> | Custom workflow | 10132 | 533 | 1185 | PXD008812 | RP |
| Yeast DeepRT<sup>52</sup> | DeepRT<sup>11</sup> | 4867 | 256 | 569 |  | RP |
| ProteomeTools PTM<sup>41</sup> | Custom workflow<br>ProteomeTools<br>PTM | 3659 | 192 | 428 | PXD009449 | RP |
| Plasma lumos 2h<sup>47</sup> | Custom workflow | 2997 | 157 | 350 | PXD013477 | RP |
| HeLa DeepRT<sup>53</sup> | DeepRT<sup>11</sup> | 2917 | 153 | 341 |  | RP |

**Supplemental Table 3:**
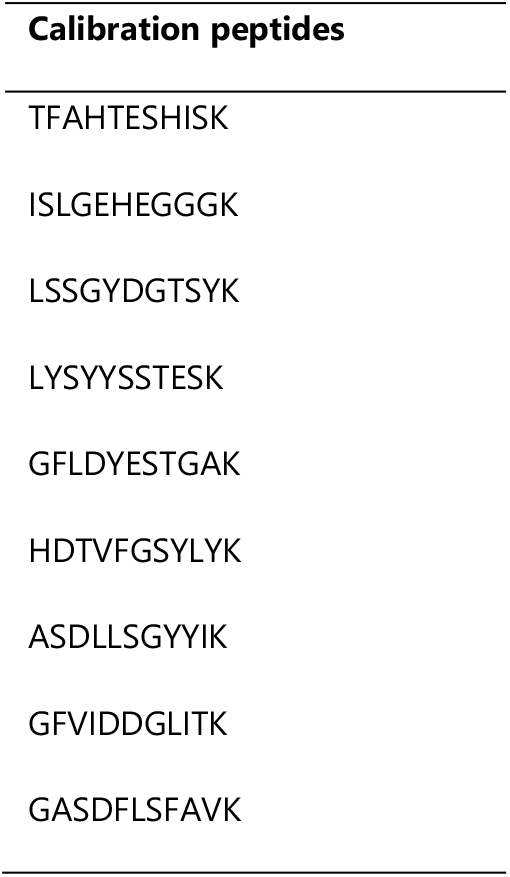
Peptide used to calibrate retention time in the *Custom workflow ProteomeTools*

**Supplemental Table 4:**
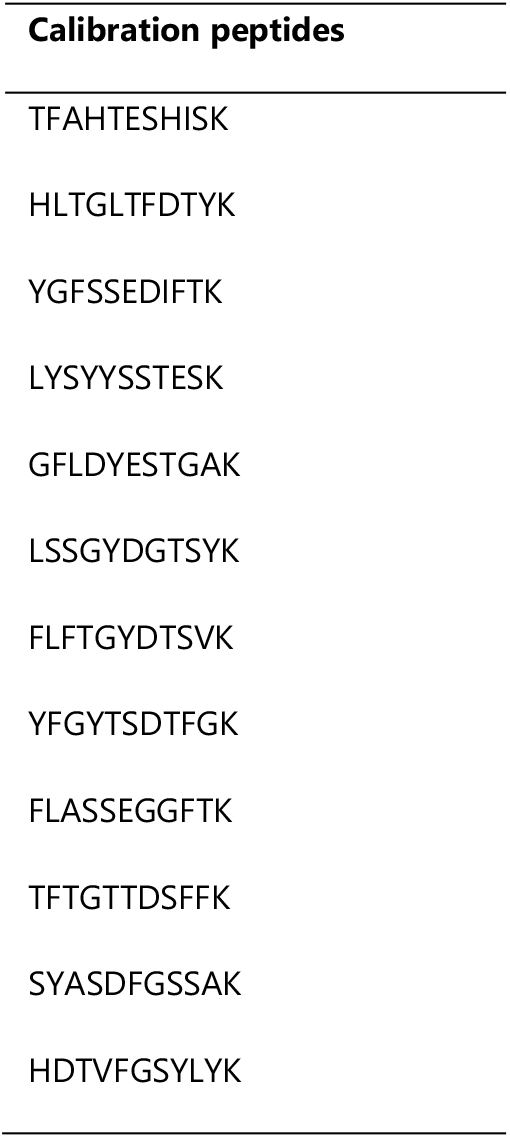
Peptide used to calibrate retention time in the *Custom workflow ProteomeTools PTM*

